# Lactate-fueled hydride transfer metabolon drives breast cancer metastasis

**DOI:** 10.64898/2026.06.16.732576

**Authors:** Sebastian Igelmann, Predrag Jovanovic, Véronique Bourdeau, Nicolas Curdy, Lesley Zhan, Ranveer Palia, David Papadopoli, Daina Avizonis, Marilène Paquet, Shannon McLaughlan, Tara Nincic-Gajic, Lorne Taylor, Jean-François Trempe, Josie Ursini-Siegel, Gerardo Ferbeyre, Ivan Topisirovic

## Abstract

Metabolic plasticity of cancer cells plays a major role in metastasis. Use of alternative carbon fuels (e.g., lactate) boosts metabolic plasticity, but the underlying mechanisms remain obscure. We show that lactate dehydrogenase B (LDHB) cooperates with the hydride ion transfer complex (HTC) comprised of pyruvate carboxylase (PC), malic enzyme (ME1) and malate dehydrogenase (MDH1) to form a metabolon that confers metabolic flexibility through lactate assimilation under physiological conditions. HTC/LDHB metabolon assembles in the aggressive breast cancer subtypes and reprograms nicotinamide adenine dinucleotide (NAD) metabolism to promote migration, invasion and escape from anoikis - thereby driving metastasis. Altogether, this work identifies a lactate-fueled metabolon that propels metastatic dissemination of breast cancer.

## Introduction

Metastatic cascade comprises a continuum of steps whereby a small subset of cancer cells develops the ability to migrate and invade neighboring tissues, extravasate, survive in lymph and/or blood circulation, colonize distant tissues and outgrow secondary tumors (Fares *et al*, 2020). To metastasize, cancer cells must overcome a variety of diverse stressors associated with distinct steps of metastatic cascade (Luzzi *et al*, 1998; Weiss, 1990). Metabolic rewiring is a hallmark of neoplasia (Vander Heiden & DeBerardinis, 2017; Zhu & Thompson, 2019) and it plays a prominent role in adaptive stress responses allowing cancer progression and dissemination (Bergers & Fendt, 2021; Faubert *et al*, 2020). Emerging evidence suggests that cancer cells can dynamically remodel their metabolism to adapt to their constantly changing environment (Biondini *et al*, 2024; Hulea *et al*, 2018; Lehuede *et al*, 2016; McGuirk *et al*, 2020). Notwithstanding that metabolic plasticity of cancer cells is thought to play a major role in metastatic spread of cancer cells, the underpinning mechanisms are still unclear.

The maintenance of redox homeostasis is a key component of metabolic adaptations of cancer cells (Nakamura & Takada, 2021; Tasdogan *et al*, 2021). The nicotinamide adenine dinucleotide (NAD) and NAD phosphate (NADP) are key cofactors that fuel metabolic reactions and detoxify reactive oxygen species (ROS) (Chiarugi *et al*, 2012; Chini *et al*, 2021). Oxidized NAD (NAD^+^) is critical for numerous metabolic pathways including glycolysis and aspartate synthesis, which in turn are required for the production of non-essential amino acids and nucleotides (Birsoy *et al*, 2015; Sullivan *et al*, 2015). Conversely, reduced NADP (NADPH) contributes to the reductive biosynthetic pathways (e.g., lipogenesis) and plays a pivotal role in ROS scavenging (Berkholz *et al*, 2008; Zakim & Herman, 1969).

We have previously identified a cytoplasmic complex which reprograms NAD metabolism in cells escaping senescence (Igelmann *et al*, 2021). Formed by pyruvate carboxylase (PC), malate dehydrogenase 1 (MDH1), and malic enzyme 1 (ME1), this hydride transfer complex (HTC) catalyzes a cyclic reaction wherein PC converts pyruvate to oxaloacetate (OAA), which is then used by MDH1 to generate malate and produce NAD^+^ from NADH (Igelmann *et al*., 2021). Finally, malate is converted back to pyruvate by ME1, closing the cycle while producing NADPH from NADP^+^ (Igelmann *et al*., 2021). Altogether, HTC cycle transfers, under physiological conditions, hydride ions from NADH to NADP^+^ to produce NAD^+^ and NADPH which allows cells to circumvent oncogene-induced senescence (Igelmann *et al*., 2021), which is a major tumor-suppressive mechanism (Wiley & Campisi, 2016).

Recently, it was demonstrated that cancer cells utilize lactate to fuel their metabolism (Faubert *et al*, 2017; Hui *et al*, 2017). Herein, we found that lactate dehydrogenase B (LDHB) associates with HTC to form a previously unrecognized HTC/LDHB metabolon. LDHB utilizes lactate to feed HTC thus driving breast cancer growth and dramatically increasing metastatic dissemination *in vivo*. These findings highlight a hitherto unrecognized, physiologically relevant mechanism of cancer cell plasticity through which lactate is used to maintain redox metabolism thus sustaining tumor progression and metastatic dissemination.

## Results

### HTC assembles in breast cancer but not in non-malignant cells and patient tissues

We have previously demonstrated that HTC strongly increased the ability of cells to overcome oncogene-induced senescence (Igelmann *et al*., 2021). To investigate the potential role of HTC in cancer, we used the proximity ligation assay (PLA) to quantify HTC foci in a panel of breast cancer cell lines representative of different histological subtypes. Here, HTC foci formed under physiological conditions and were found to be 5-10 fold more abundant in triple-negative breast cancer (TNBC) cells (BT-20, MDA-MB-231, MDA-MB-468) and HER2+ subtypes (BT-474, AU565) relative to ER+/HER2- (MCF-7, T47D) breast cancer cells (Fig. 1A, B). In turn, HTC foci were not readily detectable in non-transformed MCF10A breast epithelial cells (Fig. 1A, B). Intriguingly, differences in the number of HTC foci between breast cancer cell lines did not appear to be directly influenced by the expression levels of its subunits, PC, MDH1, and ME1 (Fig. EV1A). Importantly, PLA on human breast cancer patient samples revealed that HTC formation was the highest in TNBC tissue samples, intermediate in HER2+, lowest in ER+/HER2-and absent in normal breast tissue (Fig. 1C, D). Taking advantage of isogenic murine breast cancer models, we next quantified HTC foci in the transformed NMuMG-NT model (Ursini-Siegel *et al*, 2007) versus the non transformed NMuMG counterpart. HTC foci formed in ∼30% of transformed NMuMG-NT breast cancer cells while being largely absent in their non-transformed NMuMG counterparts (Fig. 1E, F). Together, these results show that HTC selectively forms in breast cancers but not in non-malignant cells. Moreover, the number of HTC foci appeared to increase proportionally with tumor aggressiveness as higher HTC foci levels were observed in TNBC and HER2+ tumors under physiological conditions, two subtypes associated with poor outcomes, while being the lowest in less aggressive ER+/HER2- breast cancers (Fig. 1A, C).

**Figure 1.**
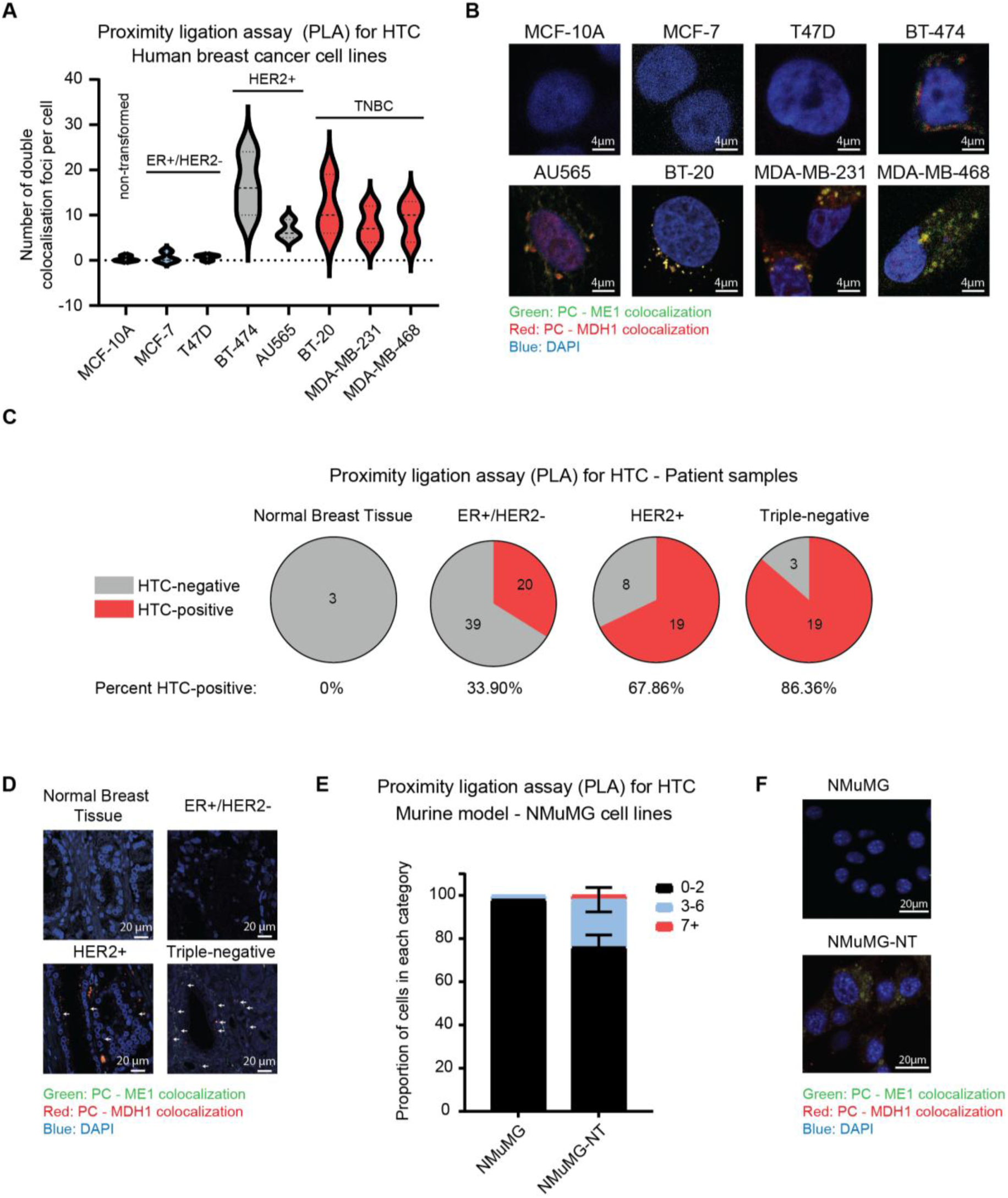
HTC foci mark aggressive breast cancer subtypes. **(A)** HTC foci were quantified using double proximity ligation assay (PLA) in a panel of non-transformed and breast cancer cell lines of indicated molecular subtypes. **(B)** Representative images of quantifications presented in panel A. **(C)** HTC foci were quantified via PLA in human patient samples belonging to indicated molecular subtypes. **(D)** Representative images of quantifications presented in panel C. **(E)** PLA was used to quantify HTC foci in a matched pair of non-transformed (NMuMG) and cancerous (NMuMG-NT) murine breast epithelial cells. **(F)** Representative images of quantifications presented in panel E. **(A-F)** All cell culture experiments were performed in three independent replicates, while five high-power fields (HPFs) per PLA replicate were quantified.

### HTC drives proliferation of breast cancer but not non-malignant cells

To investigate the functional relevance of HTC assembly in breast cancer cells, we monitored the effects of depleting HTC subunits (i.e., MDH1, ME1 or PC) on cell proliferation across different breast cancer cell lines and non-transformed cells (Fig. EV1B-G). Representative cell lines of each breast cancer subtype were employed including T47D (ER+/HER2-), BT-474 (HER2+), MDA-MB-231 and MDA-MB-468 (TNBC) cells, along with the immortalized but non-transformed control MCF-10A cells. Depletion of either of the HTC subunits only modestly decreased proliferation of MCF-10A and T47D cells (∼20% reduction compared to controls; Fig. EV1B, C, G). MCF-10A and T47D cells were previously found to be devoid or have low HTC foci levels, respectively (Fig. 1A, B). In contrast, depletion of each HTC subunit dramatically reduced proliferation of BT-474, MDA-MB-231 and MDA-MB-468 cells (i.e., ∼65% reduction compared to controls) (Fig. EV1D-G), all of which exhibit high HTC foci levels (Fig. 1A, B). These data suggest that HTC function is selectively required to drive proliferation of HER2+ and triple-negative breast cancer cells which are the subtypes presenting with the highest aggressiveness and HTC foci levels.

**Figure EV1.**
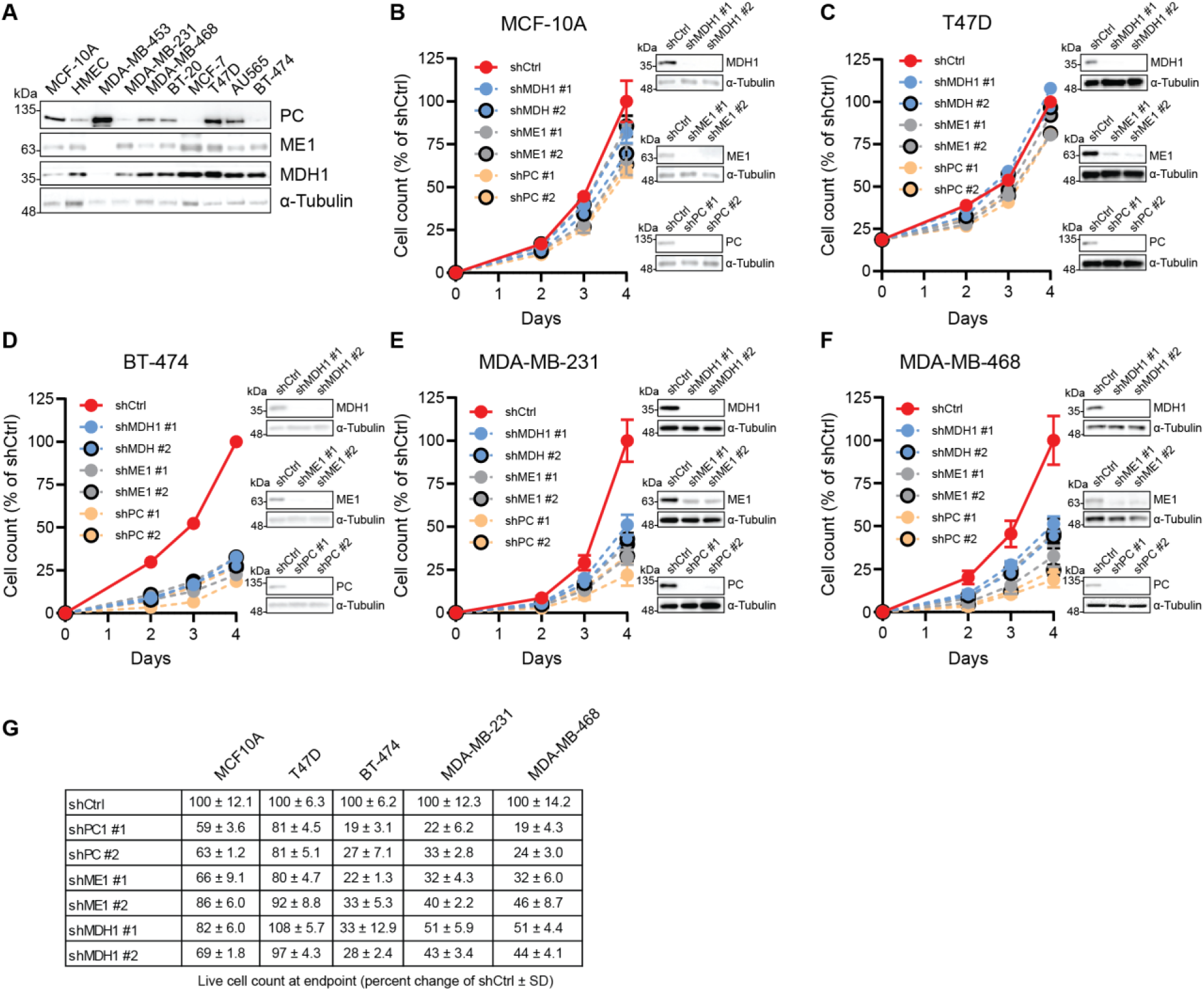
HTC drives proliferation of aggressive breast cancer cell lines. **(A-F)** Levels of PC, ME1 and MDH1 subunits of HTC were monitored by immunoblotting using indicated antibodies across the panel of breast cancer cells lines and in indicated cells in which PC, MDH1 and ME1 were depleted by shRNA. α-Tubulin served as a loading control**. (B-F)** Cell proliferation in designated control scrambled shRNA (shCtrl) expressing cells or cell lines in which PC, ME1 or MDH1 were depleted by two independent shRNAs was monitored by cell counting with trypan blue exclusion over indicated time periods. **(G)** Quantifications with standard deviations (SD) of cell proliferation from panels B-F at day 4 are shown as percent change compared to scrambled shRNA controls (shCtrl). All cell proliferation experiments were done in three independent replicates.

### HTC promotes breast tumor growth

We next determined whether HTC is implicated in breast cancer growth *in vivo*. To this end, we overexpressed PC, MDH1 and ME1 (referred to as “HTC”) in the NMuMG-NT (HER2+) cells via lentiviral infection (see methods; Fig. 2A). Cells were then orthotopically injected into the mammary fat pad of SCID-Beige mice. A ∼2 fold increase in the growth rate of HTC overexpressing tumors was observed relative to those harboring control vectors (Fig. 2B). Additionally, HTC overexpression was also established in the 67NR cells (non-aggressive cells derived from the TNBC 4T1 model) (Dexter *et al*, 1978; Heppner *et al*, 1978) (Fig. 2C), followed by orthotopic injection into the mammary fat pad of BALB/c mice (Fig. 2D). Here, we observed a ∼1.5 fold increase in the tumor growth rate of HTC overexpressing tumors compared to those infected with control vectors (Fig. 2D). Moreover, simultaneous CRISPR-Cas9-mediated abrogation of PC, MDH1 and ME1 expression (referred to as “HTC KO”) in 4T1-526 cells (aggressive cells derived from the TNBC 4T1 model) (Rose *et al*, 2010) (Fig. 2E), significantly impaired tumor engraftment and abolished tumor growth following orthotopic injection into the mammary fat pad of BALB/c mice (Fig. 2F). This was attributable to HTC subunit expression, and not off-target effects of gene deletion, as tumor growth was re-established by restoring expression of HTC subunits in the HTC KO 4T1-526 cells (indicated as “HTC Rescue”) (Fig. 2E, F). These results demonstrate that HTC promotes breast tumor growth *in vivo* in multiple murine models.

**Figure 2.**
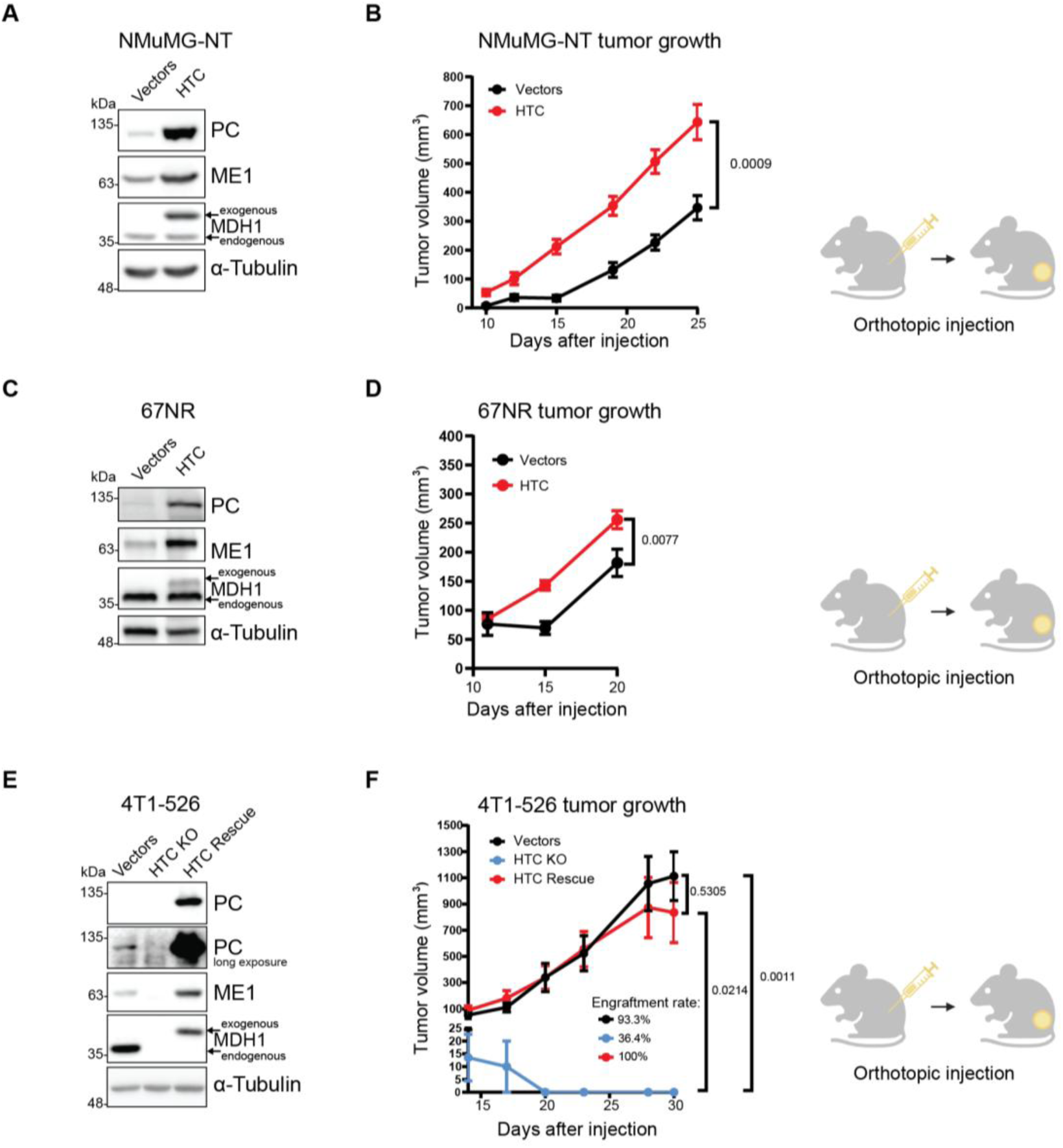
HTC promotes breast tumor growth. **(A)** Levels of indicated proteins in NMuMG-NT vector control or HTC overexpressing cells were determined by immunoblotting. α-Tubulin served as a loading control. **(B)** Control (Vectors-black, n=10) or PC/MDH1/ME1 overexpressing (HTC-red, n=10) NMuMG-NT breast cancer cells were orthotopically injected into a mammary fat pad of SCID-Beige mice. **(C)** Levels of indicated proteins in 67NR vector control or HTC overexpressing cells were determined by western blotting. α-Tubulin served as a loading control. **(D)** Control (Vectors-black, n=20) or PC/MDH1/ME1 overexpressing (HTC-red, n=20) 67NR breast cancer cells were orthotopically injected into a mammary fat pad of BALB/c mice. **(E)** Levels of indicated proteins in 4T1-526 vector control, PC/MDH1/ME1 KO (“HTC KO”) or HTC KO cells re-expressing PC/MDH1/ME1 (“HTC Rescue”) were determined by immunoblotting. α-Tubulin served as a loading control. **(F)** Control (Vectors-black, n=15), PC/MDH1/ME1 KO (HTC KO-blue, n=16) or PC/MDH1/ME1 KO 4T1-526 breast cancer cells in which PC/MDH1/ME1 were re-expressed (HTC Rescue-red, n=15) were orthotopically injected into a mammary fat pad of BALB/c mice. **(B, D, F)** Tumor volume at indicated times was monitored by caliper measurements and presented as mean and standard error of the mean (SEM) values. P-values are indicated. Statistical analysis was performed using unpaired t-test **(B, D)**, or One-way ANOVA **(F)**.

### HTC promotes breast cancer metastasis

Considering that metabolic reprogramming plays a major role in adaptation to stress across the metastatic cascade (Faubert *et al*., 2020), we next investigated whether HTC also contributes to breast cancer metastasis. To test this, we quantified lung metastases formed after orthotopic injection of NMuMG-NT cells (i.e., spontaneous metastasis). This approach recapitulates the entire sequence of the metastatic cascade. We found that HTC overexpression substantially increased the number of mice with spontaneous lung metastases compared to control tumors (80% HTC overexpression vs 50% vector controls) (Fig. 3A). Moreover, we observed a ∼5 fold increase in lung metastatic burden in mice bearing HTC-overexpressing tumors compared to controls, despite resections taking place at equal primary tumor volumes (Fig. 3B).

**Figure 3.**
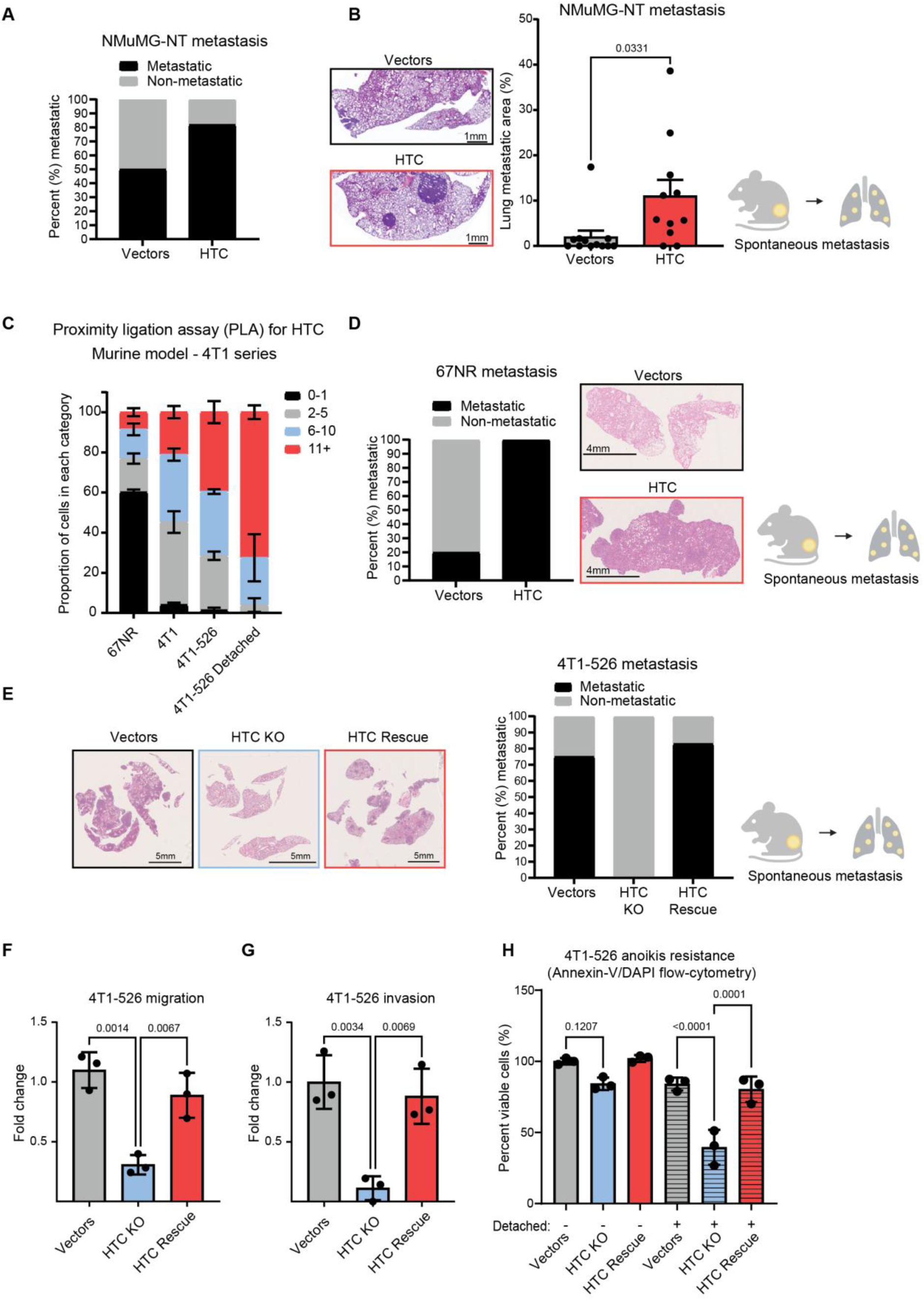
HTC promotes breast cancer metastasis. **(A)** Proportions of animals harboring metastases were determined by hematoxylin and eosin (H&E) staining and scanning of lung sections from SCID-Beige mice injected orthotopically with PC/MDH1/ME1 overexpressing NMuMG-NT breast cancer cells (Vectors n=12; HTC n=11). Lungs were collected 21 days after resecting tumors at a volume of 300 mm^3^. **(B)** Percentage of the areas of the lungs covered by metastatic lesions were quantified through ImageScope after scanning of H&E-stained lung sections in animals from panel A. Bars represent SEM values. P-values are indicated (unpaired t-test). **(C)** Double PLA was used to quantify HTC foci in the 4T1 series of murine breast cancer cell lines of varying metastatic potential. Five high-power fields (HPFs) per replicate were quantified. **(D)** Proportions of animals with lung metastases in a BALB/c cohort injected orthotopically with 67NR breast cancer cells overexpressing PC/MDH1/ME1 (HTC, n=10), or control vectors (Vectors, n=10) were determined upon scanning of H&E-stained slides, upon necropsy of animals when tumor volumes reached 1000 mm^3^. (**E**) Proportions of animals harboring lung metastases were determined upon scanning of H&E-stained lung sections in Control (Vectors, n=12), PC/MDH1/ME1 KO (HTC KO, n=16) or PC/MDH1/ME1 KO 4T1-526 breast cancer cells in which PC/MDH1/ME1 were re-expressed (HTC Rescue, n=12), upon necropsy of animals when Vectors and HTC Rescue tumor volumes reached 1000 mm^3^. **(F)** Migration, **(G)** invasion or **(H)** anoikis were measured in control (Vectors-gray), PC/MDH1/ME1 KO (HTC KO-blue) or PC/MDH1/ME1 KO 4T1-526 breast cancer cells in which PC/MDH1/ME1 were re-expressed (HTC Rescue-red) 24 h after seeding on to (**F**) uncoated and **(G)** matrigel-coated transwell inserts or **(H)** by using Annexin-V/DAPI staining coupled with flow-cytometry 24 h after cell detachment. **(F, G)** migratory and invasive cells that traversed the insert were counted upon formalin fixing and crystal violet staining to reveal fold change differences. Each bar graph shows mean and SD values of experiments performed in three independent replicates, where p-values are indicated (One-way ANOVA).

Next, we used the 4T1 series of breast cancer cell lines - all derived from a single spontaneously arising mammary tumor of BALB/cfC3H mice but exhibiting varying levels of metastatic proficiency (Dexter *et al*., 1978; Heppner *et al*., 1978; Rose *et al*., 2010). Intriguingly, the numbers of HTC foci, determined by PLA, were proportional to the metastatic ability, as the poorly metastatic 67NR cells showed lowest HTC foci levels, parental 4T1 exhibited intermediate HTC foci abundance, while HTC foci levels were the highest in the highly metastatic 4T1-526 cells (Fig. 3C). Notably, metastasis-relevant stress (cell detachment) further bolstered HTC foci levels in 4T1-526 cells (Fig. 3C).

Remarkably, HTC overexpression in poorly metastatic 67NR cells strongly increased their lung metastatic potential (Fig. 3D). Herein, HTC overexpressing 67NR cells formed metastasis in 100% of the mice as compared to only 20% of mice harboring metastasis in the 67NR vector control group (Fig 3D). Of note, necropsies were performed at equal primary tumor volumes in all animals. This indicates that HTC stimulates metastatic dissemination of breast cancer cells independently of its effects on primary tumor growth. As expected, highly metastatic 4T1-526 HTC KO breast cancer cells did not form metastasis due to impeded primary tumor formation (Fig. 2F, 3E). Re-expressing HTC subunits in 4T1-526 HTC KO breast cancer cells restored their metastatic potential (Fig. 3E). Overall, these findings indicate that HTC is required for the metastatic spread of breast cancer cells *in vivo*.

We next set out to identify the step(s) in the metastatic cascade during which HTC plays a critical role using the highly metastatic 4T1-526 breast cancer cell line. HTC abrogation in 4T1-526 HTC KO cells reduced transwell migration and invasion as compared to HTC proficient cells (Fig. 3F, G). Although the HTC status of cancer cells did not exert a major effect on cell viability under adherent conditions, abrogation of the HTC components significantly decreased survival of detached cells, thus indicating that HTC enables breast cancer cells to overcome anoikis (Fig. 3H). Taken together, these results demonstrate that HTC promotes the early steps of the metastatic cascade to enhance breast cancer metastasis *in vivo*.

### Lactate fuels the HTC cycle

We next sought to delineate the mechanisms underpinning the pro-metastatic function of HTC. To achieve this, we set out to identify the potential HTC interacting factors using ME1 immunoprecipitation followed by mass spectrometry (IP-MS). Initial studies were carried out in prostate cancer PC-3 cells, which we have previously demonstrated exhibit high numbers of HTC foci (Igelmann *et al*., 2021). As expected, MDH1 and PC were among the most abundant proteins that co-immunoprecipitated with ME1 (Fig. EV2A-B). In addition, we identified LDHB as significantly enriched in ME-1 immunoprecipitates (Fig. EV2A-B), with spectral counts in line with previous reports in the literature (Goncalves *et al*, 2022; Moghimi *et al*, 2022; Monaco *et al*, 2024). LDHB produces pyruvate and NADH from lactate and NAD^+^ (Farhana & Lappin, 2025), both of which are utilized by HTC (Igelmann *et al*., 2021). The opposite reaction is catalyzed by LDHA (Farhana & Lappin, 2025). Since emerging results suggest that lactate serves as a major carbon source for cancer cells (Faubert *et al*., 2017; Hui *et al*., 2017), we hypothesized that LDHB may metabolize lactate to fuel the HTC cycle with pyruvate and NADH. To validate the IP-MS results, we performed anti-ME1 immunoprecipitation in MDA-MB-231 (Fig. 4A) and MDA-MB-468 (Fig. 4B) human TNBC cells and confirmed that LDHB associates with HTC. Next, we determined whether HTC promotes lactate utilization by breast tumors under physiological conditions *in vivo* using metabolic tracing of deuterium labelled lactate (Fig. 4C). Herein, we orthotopically injected HTC overexpressing or vector control NMuMG-NT cells and resected tumors when volumes reached 300 mm^3^ (Fig. 4D). Tumors were then sectioned using a vibratome and cultured in media supplemented with deuterium labelled lactate (Fig. 4D). Gas chromatography-mass spectrometry (GC-MS) analysis revealed that HTC overexpressing NMuMG-NT tumors had increased labeling of malate (m+1) with lactate-derived deuterium compared to vector control tumors, thus suggesting that HTC increases lactate utilization in breast tumors (Fig. 4E). During conversion of lactate to pyruvate by LDHB the produced NADH accepts the deuterium isotope from the labelled lactate, resulting in label-free pyruvate (Fig. 4C). By consecutive action of MDH1 and ME1, the deuterium label is then transferred from NADH to malate (m+1), and finally from malate to NADPH (Fig. 4C). To confirm that the lactate-derived hydride ion transfer is indeed catalyzed by HTC, we established expression of the R132H mutant of cytosolic isocitrate dehydrogenase 1 (IDH1) which has impaired affinity for isocitrate and instead catalyzes reduction of a-ketoglutarate to 2-hydroxyglutarate (2-HG) while converting NADPH to NADP^+^ (Dang *et al*, 2009; Yan *et al*, 2009; Zhao *et al*, 2009) (Fig. 4F, Fig. EV2C). Deuterium from lactate-derived NADPH is incorporated into 2-HG by the cytosolic R132H IDH1 mutant (Fig. 4F). Therefore, to test whether lactate feeds the HTC cycle in the cytosol, we labelled control, HTC KO and HTC-rescued 4T1-526 cells (Fig. 2E) expressing the R132H IDH1 mutant (Fig. EV2C) with deuterated lactate (Fig. 4G). Deuterium-labelled lactate was metabolized into malate (m+1) only in HTC proficient but not HTC deficient cells (Fig. 4H). As expected, pyruvate labelling was negligible across all cell lines (Fig. 4I). Crucially, incorporation of lactate-derived deuterium into 2-HG (m+1) was markedly higher in HTC proficient vs. deficient cells (Fig. 4J). Conversely, a-ketoglutarate m+1 labelling was low in all cell lines (Fig. 4K), which is consistent with the gain of function of the R132H IDH1 mutant. Collectively these results demonstrate that lactate fuels the HTC cycle.

**Figure 4.**
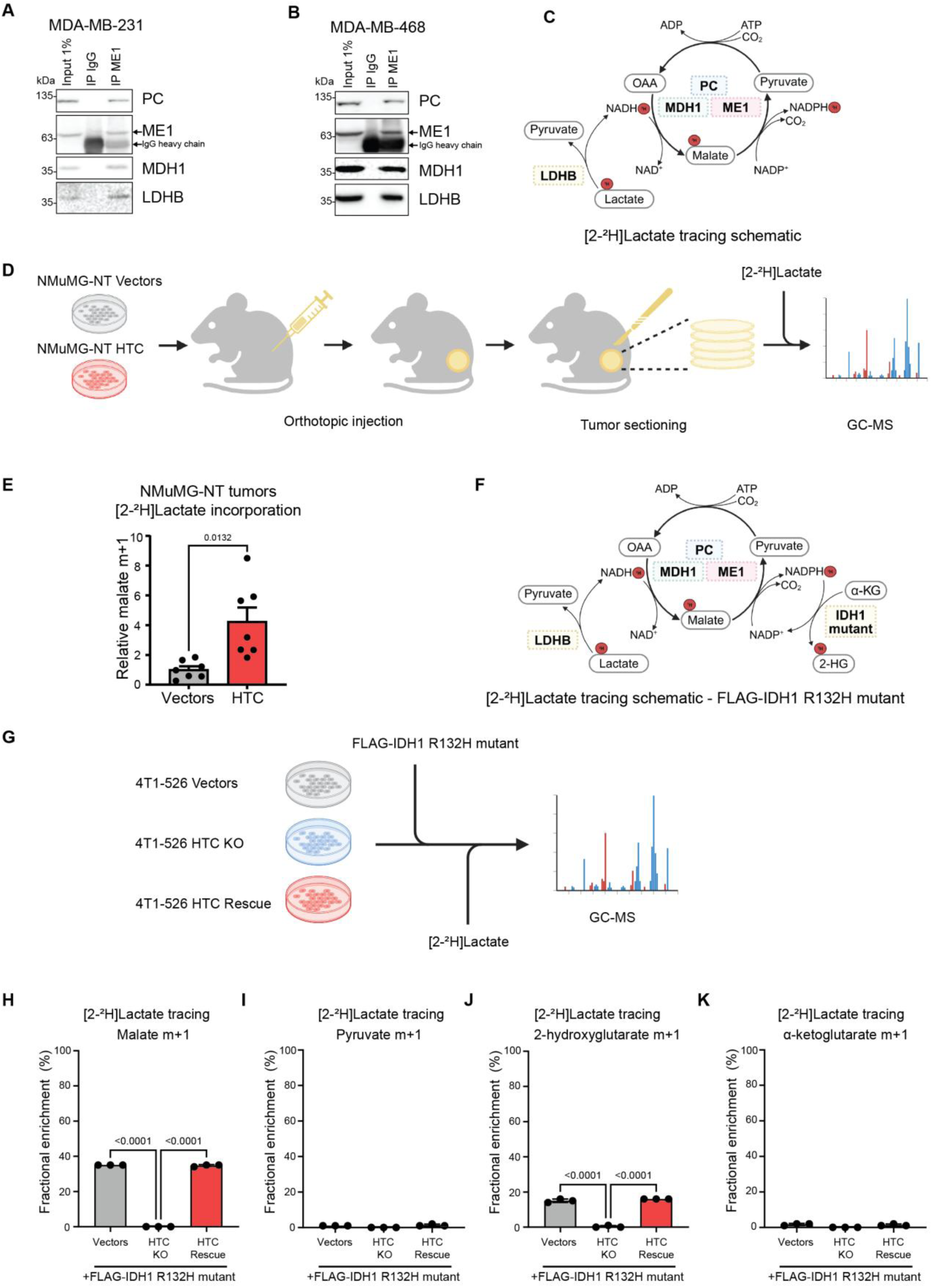
HTC is fueled by lactate via LDHB. Levels of indicated proteins in input (1%), control (IgG) or anti-ME1 antibody co-immunoprecipitates were monitored by immunoblotting in MDA-MB-231 **(A)** and MDA-MB-468 **(B)** human breast cancer cells. **(C)** Schematic depicting predicted labelling of metabolites by HTC using [2-^2^H]-lactate. **(D)** Schematic of deuterium-labelled lactate tracing in live breast tumor sections. **(E)** Incorporation of [^2^H] from [2-^2^H]-labelled lactate into malate was quantified using Gas Chromatography-Mass Spectrometry (GC-MS) in sections of PC/MDH1/ME1 overexpressing (HTC-red, n=7), or control vector expressing (Vectors-black, n=7) NMuMG-NT tumors. Tumors were resected from SCID-Beige mice at volumes of 300 mm^3^ and sectioned using a vibratome. Mean, SEM and p-values are shown (unpaired t-test). **(F)** Schematic depicting predicted labelling of metabolites by HTC and R132H IDH1 mutant using [2-^2^H]-lactate. **(G)** Schematic and **(H-K)** metabolic tracing of [2-^2^H]-lactate by GC-MS in R132H IDH1 mutant-expressing or vector control (Vectors-gray), PC/MDH1/ME1 KO (HTC KO-blue) or PC/MDH1/ME1 KO 4T1-526 breast cancer cells in which PC/MDH1/ME1 were re-expressed (HTC Rescue-red). Mean, SD and p-values of three independent replicates are shown (one-way ANOVA).

**Figure EV2.**
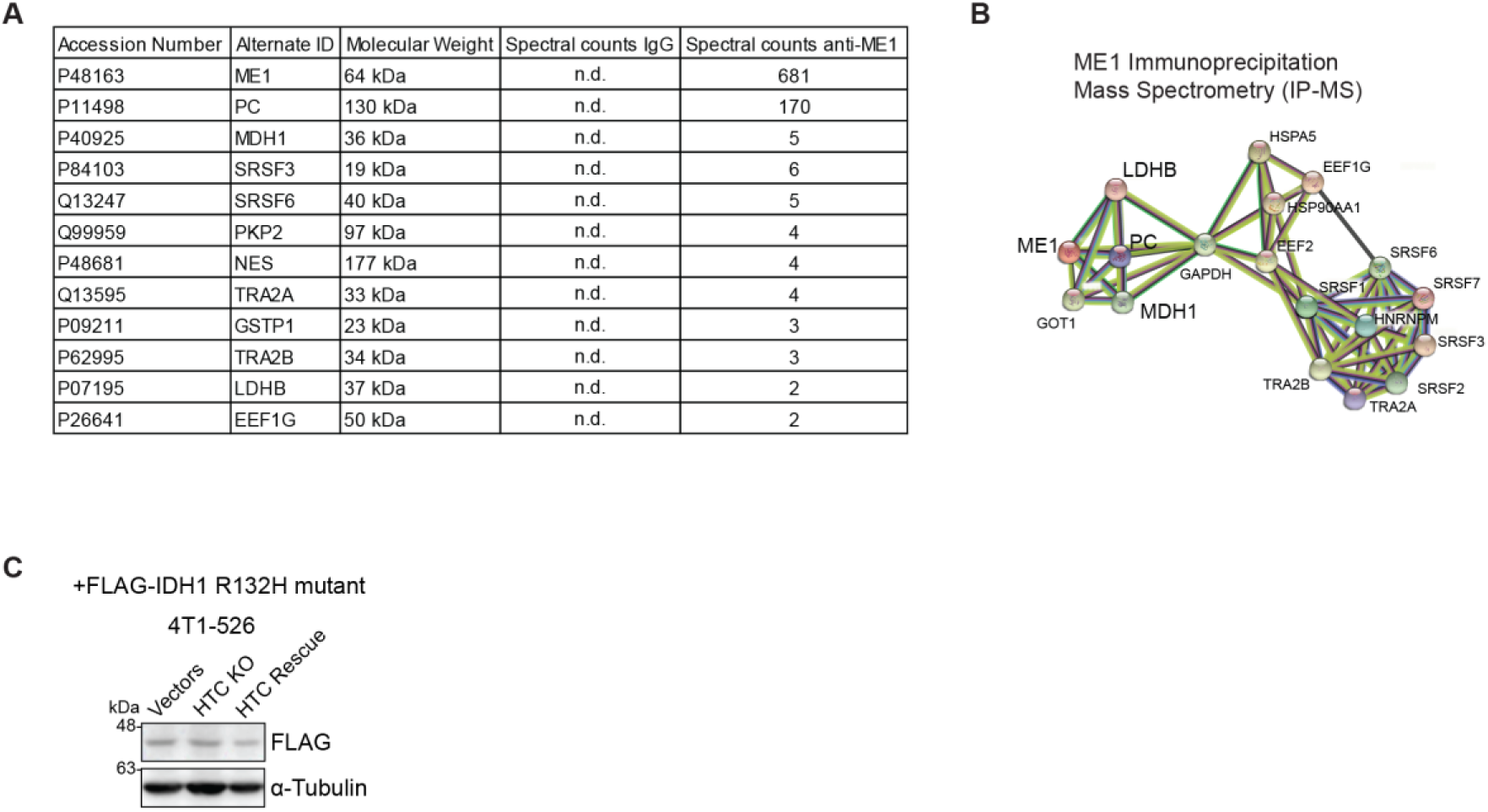
HTC interactome. **(A)** Anti-ME1 antibody immunoprecipitation was followed by high-performance liquid chromatography (HPLC)-mass spectrometry to identify potential interactors of ME1 in PC-3 prostate cancer cells. Identified candidate proteins and respective spectral counts are shown. **(B)** The top ranked identified proteins were analyzed using the STRING analysis to map potential interactions, indicated as lines in the plot. **(C)** Immunoblotting was used to validate FLAG-tagged IDH1 R132H mutant expression in Control (Vectors), PC/MDH1/ME1 KO (HTC KO) or PC/MDH1/ME1 KO 4T1-526 breast cancer cells in which PC/MDH1/ME1 were re-expressed (HTC-Rescue). α-Tubulin served as a loading control.

### HTC/LDHB lactate-fueled metabolon drives breast cancer metastasis

We next set out to determine whether LDHB plays a role in pro-tumorigenic and pro-metastatic effects of HTC. Herein, we depleted LDHB or LDHA (which was not found to be associated with HTC in the IP-MS experiment) in 4T1 cells overexpressing HTC or control vectors (Fig. 5A). In contrast to human cells (Fig. 4A, B), immunoblotting of murine 4T1 cells using the LDHB antibody revealed two bands migrating near the expected 35 kDa size. The slower migrating band was determined to be LDHB whereas the faster migrating one corresponded to LDHA, as judged by the reduced band intensity upon shRNA-mediated depletion of LDHB or LDHA, respectively (Fig. 5A). Of note, two separate shRNA constructs were used to deplete expression of each gene of interest. We selected “shLDHB #2” for *in vivo* studies as it led to the greatest reduction of LDHB protein levels.

**Figure 5.**
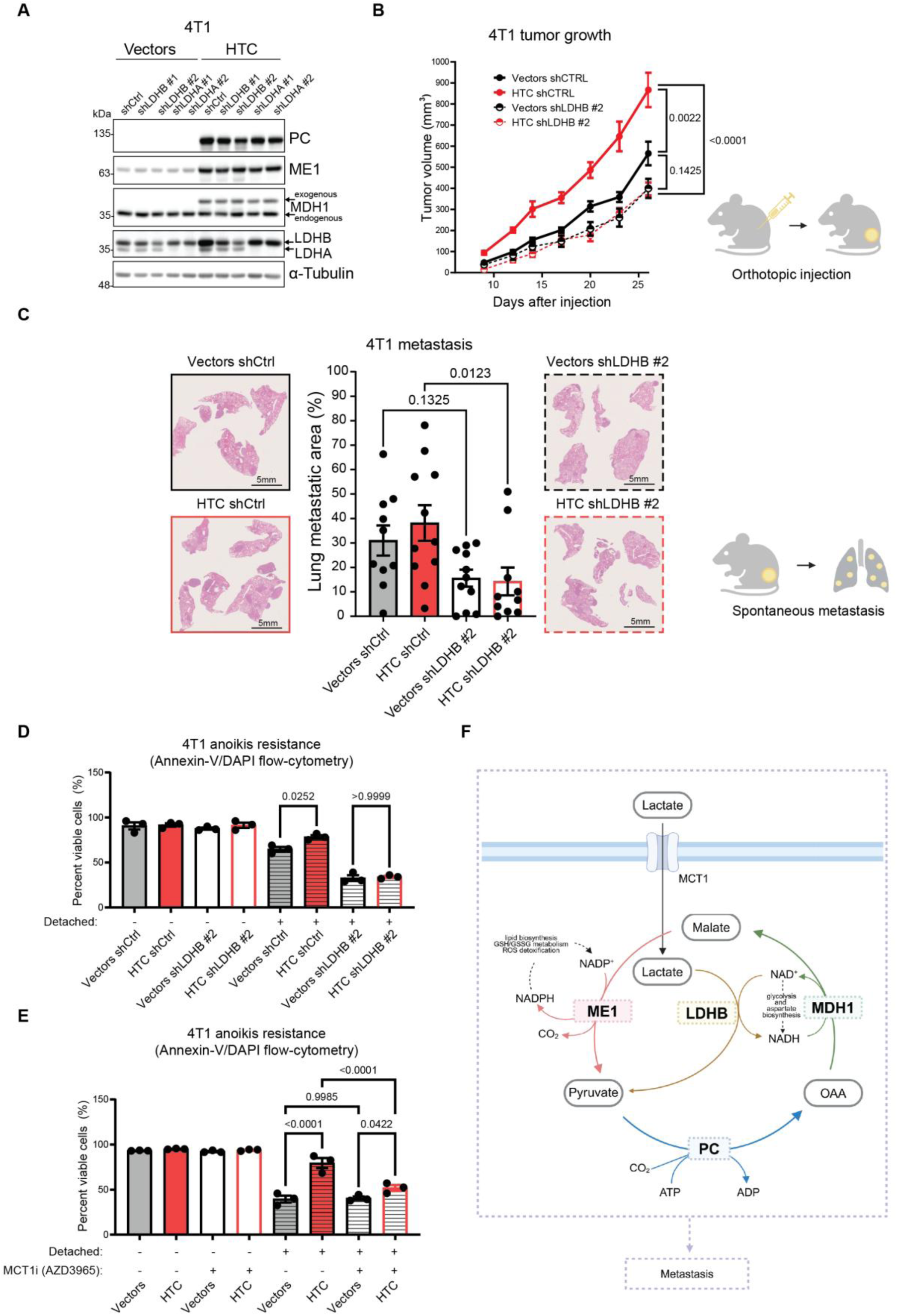
HTC/LDHB metabolon drives breast cancer metastasis. **(A)** 4T1 cells expressing vector controls (Vectors) or overexpressing PC/MDH1/ME1 (HTC) were transduced with scrambled control shRNA (shCtrl) or two independent LDHA- or LDHB-targeting shRNAs. Levels of indicated proteins were determined by immunoblotting. α-Tubulin served as a loading control. **(B)** Tumor growth was measured using calipers in BALB/c mice following mammary fat pad injection of 4T1 breast cancer cells expressing vector control (Vectors-black), or overexpressing PC/MDH1/ME1 (HTC-red) that were infected by either scrambled control shRNA (shCtrl-full line) or LDHB-targeting shRNA (shLDHB #2-broken line), producing the indicated four groups (Vectors shCtrl, n=10; HTC shCtrl, n=11, Vectors shLDHB #2, n=11, HTC shLDHB #2, n=10). Mean, SEM and p-values are shown (One-way ANOVA). **(C)** Lung metastases were quantified by scanning of H&E-stained lung sections using ImageScope in each group from panel A. Mice were necropsied when respective tumor volumes reached 1000 mm^3^. Mean, SEM and p-values are shown, while statistical comparison was done using One-way ANOVA. **(D-E)** Anoikis was measured 24 h after cell detachment by using Annexin-V/DAPI staining coupled with flow-cytometry in cells expressing vector controls (Vectors) or overexpressing PC/MDH1/ME1 (HTC) that were either **(D)** transduced with scrambled control shRNA (shCtrl) or LDHB-targeting shRNA (shLDHB #2) or **(E)** treated with the MCT1 inhibitor (AZD3965, 10 nM) or a vehicle (DMSO). All experiments were done in three independent replicates. Mean, SD and p-values are shown (One-way ANOVA). **(F)** Schematic depicting proposed model of the study.

To determine the functional importance of LDHB in HTC-driven tumor growth and metastasis, we performed *in vivo* tumor growth and metastasis experiments with 4T1 vector control or HTC subunit overexpressing cells transduced with either scrambled control or an shRNA targeting LDHB. As previously observed (Fig. 2A-D), HTC overexpression accelerated breast cancer growth by ∼1.5 fold relative to vector control 4T1 cells upon orthotopic injection of the 4T1 cells into a mammary fat pad of BALB/c mice (Fig. 5B). Strikingly, LDHB depletion in HTC overexpressing 4T1 cells abolished the growth advantage conferred by HTC overexpression (Fig. 5B). Importantly, this was also reflected in spontaneous metastasis to the lungs where LDHB depletion significantly reduced metastasis of HTC overexpressing but not vector control 4T1 cells (Fig. 5C).

Next, we asked whether LDHB is required for pro-metastatic phenotypes conferred by HTC. Herein, we measured survival upon detachment of vector control or HTC overexpressing 4T1 breast cancer cells transduced with scrambled control (shCtrl) or shRNA targeting LDHB (shLDHB #2). Cells overexpressing HTC exhibited higher viability upon detachment when compared to vector control cells (Fig. 5D). However, this survival advantage conferred by HTC was abolished by LDHB depletion (Fig. 5D). Importantly, this was specific to LDHB, as depletion of LDHA did not exert a major effect on the survival of cells upon detachment (Fig. EV3A). This demonstrated that HTC promotes anoikis resistance in an LDHB-dependent manner. Notably, LDHB, but not LDHA, depletion reduced proliferation of MDA-MB-231 cells (Fig EV3B-C) which were previously identified as being abundant in HTC foci (Fig. 1A, B).

Monocarboxylate transporter 1 (MCT1) facilitates the transport of lactate and other monocarboxylates across the cell membrane (Garcia *et al*, 1994; Halestrap, 2013). MCT1 is also implicated in metastasis(Tasdogan *et al*, 2020). To investigate whether the HTC/LDHB metabolon relies on MCT1 for lactate import, we measured anoikis resistance in 4T1 vector control or HTC overexpressing cells treated with the inhibitor of MCT1 (AZD3965) or vehicle control. While overexpression of HTC increased anoikis resistance, this survival advantage was strongly attenuated upon inhibition of MCT1 (Fig. 5E). This suggests that lactate uptake via the MCT1 transporter is required for HTC-driven resistance to anoikis.

Surprisingly, we observed that 4T1 cells overexpressing HTC subunits exhibited increased LDHB protein levels as compared to those expressing vector control (Fig. 5A). This suggested that HTC not only interacts with LDHB but may also regulate its levels. To confirm this result, we also monitored LDHB expression in vector control, HTC KO and HTC Rescue 4T1-526 cells. This revealed that HTC-proficient cells display elevated LDHB abundance compared to HTC deficient cells at both the protein (Fig. EV3D) and mRNA levels (Fig. EV3E). This pattern of HTC-dependent regulation of LDHB expression was not detected for LDHA (Fig. EV3F) thus suggesting specific regulation of LDHB expression by HTC. Taken together, these results demonstrate that HTC not only interacts with LDHB, but also potentially bolsters its expression to form a lactate-fueled metabolon that promotes breast cancer growth and metastasis (Fig. 5F).

**Figure EV3.**
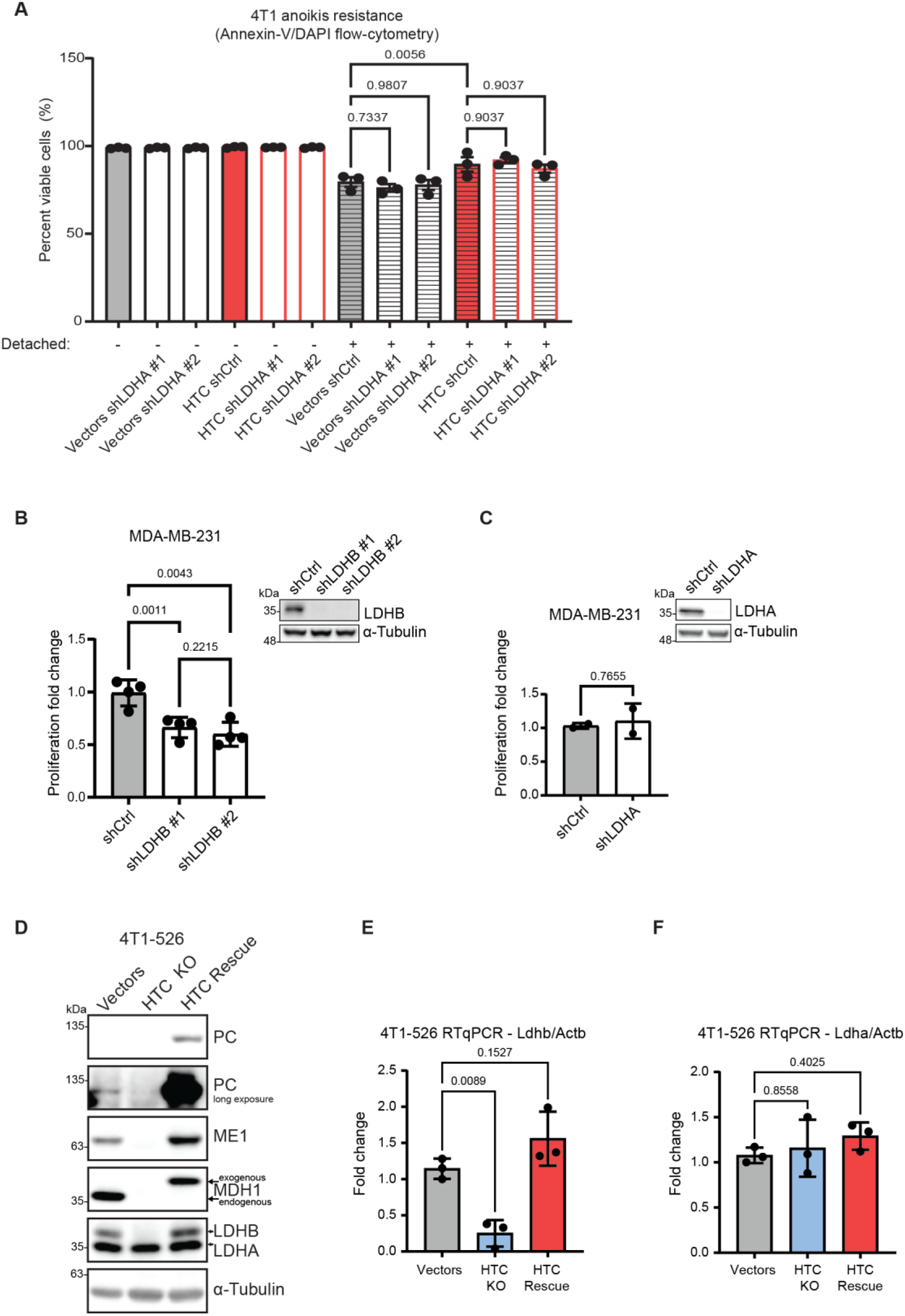
HTC co-operates with and regulates expression of LDHB, but not LDHA. **(A)** Anoikis was measured 24 h after cell detachment by Annexin-V/DAPI staining coupled with flow-cytometry in cells expressing vector controls (Vectors) or overexpressing PC/MDH1/ME1 (HTC) that were transduced with scrambled control shRNA (shCtrl) or LDHA-targeting shRNA (shLDHA #1, shLDHA #2). Experiments were done in three independent replicates wherein mean and SD values are shown. P-values are indicated (one-way ANOVA). **(B-C)** Relative cell proliferation was determined by crystal violet retention after 8 days in culture in MDA-MB-231 cells with depleted LDHB levels **(B)**, depleted LDHA levels **(C)**, or shRNA scramble control cells **(B-C)**. The experiment was done in three independent replicates whereby mean and SD values are shown. Statistical analysis was performed using one-way ANOVA (B) and unpaired t-test **(C)**, with p-values indicated. **(D)** Levels of indicated proteins in control (Vectors), PC/MDH1/ME1 KO (HTC KO) or PC/MDH1/ME1 KO 4T1-526 breast cancer cells in which PC/MDH1/ME1 were re-expressed (HTC Rescue) were monitored by immunoblotting. α-Tubulin was used as a loading control. **(E-F)** Levels of mRNA encoding LDHB **(E)** and LDHA **(F)** were determined by RT-qPCR in control (Vectors-gray), PC/MDH1/ME1 KO (HTC KO-blue) or PC/MDH1/ME1 KO 4T1-526 breast cancer cells in which PC/MDH1/ME1 were re-expressed (HTC Rescue-red). Graphs show mean and SD values of experiments performed in three independent replicates. P-values are indicated (one-way ANOVA).

## Discussion

This work uncovered HTC/LDHB metabolon that assembles under physiological conditions in aggressive breast cancers and drives tumor growth and metastasis by assimilating lactate. Recent findings show that lactate is a major carbon source that fuels cancer cells in the tumor microenvironment wherein glucose is frequently limiting (Faubert *et al*., 2017; Hui *et al*., 2017). Monocarboxylate transporter 1 (MCT1) which transports lactate across the cell membrane, has also been linked to tumor growth and metastasis (Sonveaux *et al*, 2008; Tasdogan *et al*., 2020). Emerging data demonstrate that NAD metabolism may play a major role in metastasis (Heske, 2019; Navas & Carnero, 2021; Yong *et al*, 2023). Across several cancer types, both the synthesis and redox status of NAD were found to regulate cell migration, invasion, as well as metastasis (Cheng *et al*, 2015; Santidrian *et al*, 2013; Soncini *et al*, 2014; van Horssen *et al*, 2013; Zhang *et al*, 2018). Additionally, NADPH plays a key role in ROS detoxification and biosynthetic pathways (e.g., lipogenesis) that are required for metastatic dissemination (Bergers & Fendt, 2021; Faubert *et al*., 2020; Piskounova *et al*, 2015; Wei *et al*, 2020). Our work demonstrates the previously unappreciated lactate-centered mechanism whereby HTC/LDHB metabolon utilizes lactate to regenerate pools of electron acceptors (NAD+) and reducing equivalents (NADPH) while driving growth and dissemination of neoplastic cells *in vivo*.

HTC assembly is triggered by stress. For instance, the number of HTC foci was notably higher in breast cancer cells where stress was induced by their detachment (Fig. 3C). Moreover, the number of HTC foci is induced under hypoxia (Igelmann *et al*., 2021) which is a major metastasis-promoting stressor (Jewer *et al*, 2020; Rankin & Giaccia, 2016; Schindl *et al*, 2002; Schito & Rey-Keim, 2023; Yamamoto *et al*, 2008; Yang *et al*, 2008). Collectively, these findings suggest that HTC assembly is facilitated by stressors acting across the metastatic cascade, whereby HTC cooperates with LDHB in allowing cancer cells to consume lactate.

Cancer cells exhibit flexibility in nutrient utilization and engagement of metabolic pathways. This metabolic plasticity of cancer cells facilitates adaptation to therapeutic insults and stimulates metastatic dissemination (Bergers & Fendt, 2021; Lehuede *et al*., 2016; McGuirk *et al*., 2020; Wei *et al*., 2020). Stress-induced, reversible assembly of HTC/LDHB metabolon endows cancer cells with the flexibility in nutrient utilization to regenerate NAD^+^ and NADPH. Importantly, this occurs without net carbon loss thus not depleting cellular pools of key metabolites (e.g., pyruvate). Altogether this suggests that HTC/LDHB metabolon may be a major factor underpinning metabolic plasticity of cancer cells.

HTC/LDHB metabolon assembles specifically in the aggressive breast cancer subtypes, while being largely dispensable for non-transformed cells. This alludes to a possible therapeutic window for targeting HTC/LDHB metabolon to selectively eradicate cancer cells while causing only marginal toxicity in normal tissues. Our findings thus suggest that developing therapeutic means to interfere with HTC/LDHB metabolon assembly may open new avenues to target breast cancer and/or prevent its metastatic spread in an adjuvant setting.

## Materials and Methods

### Cell lines

The cell lines used were cultured in humidified incubators at 37°C under 5% CO_2_. The respective cell lines and growth media are described below.

| Cell line | Medium | Supplementation |
| --- | --- | --- |
| MCF-7<br>T47D<br>PC-3<br>BT-474<br>AU565<br>4T1<br>4T1-526 | RPMI medium (Wisent) | 10% fetal bovine serum,<br>1% penicillin/streptomycin<br>(Wisent), and 2 mM L-glutamine |
| BT-20<br>MDA-MB-231<br>MDA-MB-468<br>MDA-MB-453<br>67NR | DMEM medium (Wisent) | 10% fetal bovine serum,<br>1% penicillin/streptomycin<br>(Wisent), and 2 mM L-glutamine |
| HMEC | EpiMax Kit for the culture of<br>mammary epithelial cell<br>lines (Wisent) | Epi Max SH: 002-013-IL,<br>Epi Max SA: 002-014-IL,<br>Epi Max SG: 002-015-TL |
| MCF-10A | DMEM/F12 (1:1) (Wisent) | 5% horse serum, 10 µg/mL<br>insulin, 0.5 µg/mL<br>hydrocortisone, 10 ng/mL<br>epidermal growth factor<br>(EGF), 100 ng/mL cholera<br>toxin, 2 mM L-glutamine |
| NMuMG | DMEM medium (Wisent) | 10% fetal bovine serum, 1% penicillin/streptomycin (Wisent), 2mM L-glutamine, 10mg/mL insulin, 10 mM HEPES |
| NMuMG-NT | DMEM medium (Wisent) | 10% fetal bovine serum, 1% penicillin/streptomycin (Wisent), 2 mM L-glutamine, 10mg/mL insulin, 10 mM HEPES, 2mg/mL puromycin |

### Proximity ligation assay (PLA)

Proximity ligation assays were done as previously described (Igelmann *et al*., 2021). Briefly, Duolink Multicolor Kit was used with red and green conjugation kits. Primary antibodies were conjugated to green and red oligos as instructed by the manufacturer, and were stored at 4°C.

To prepare for PLA, cells were fixed with 4% paraformaldehyde (PFA) in phosphate buffered saline (PBS) for 10 min. This was followed by two washes of 5 min with PBS containing 0.1 M glycine to inactivate excess PFA. Cells were then permeabilized with PBS containing 0.1 M glycine and 0.4% Triton X-100 for 5 min at 4 °C and blocked using Duolink blocking buffer for 1 h at 37 °C. Following adjustments were made to the manufacturer’s protocol. Conjugated antibodies were further diluted to 1:75 for PC-MDH1 and 1:50 for PC-ME1. They were incubated overnight at 4°C. For ligation, the multicolor ligase was diluted to 1:20, before a 30 min incubation at 37°C. Amplification and detection were done for 120 min and 45 min, respectively, both at 37°C. For wash buffer preparation, one package of wash buffer A powder was dissolved in 1 L of Milli-Q water. For wash buffer B, one package of wash buffer B powder was dissolved in 1 L of Milli-Q water. Wash buffers were stored at 4 °C for no longer than 1 month and warmed to room temperature (RT) before use. For PLA on human tissues, with following modifications: washing times, after blocking with Duolink block buffer, were increased to 10 min and done four times with wash buffer A.

Cells were washed three times for 5 min using wash buffer, wherein the first wash contained 300 ng/mL DAPI. Mounting was done using Vectorshield, while the edges were sealed off with nail polish. Mounted cells were kept at 4°C for a minimum of 24 h and were incubated in a closed slide box at RT for at least 1 h before imaging. Confocal microscope imaging was done using the Zeiss LSM 800 with a spectral analysis detector. All images were acquired sequentially and with a maximal airy unit of 2.

To quantify PLA signal colocalization, the filter set 77 of Zeiss was used. As such, the colocalization appears as yellow, while no colocalization is seen as either red or green in the oculars. For quantification, 15 random foci per condition were chosen and analyzed in ImageJ and Fiji.

### Cell harvesting, protein quantification, SDS-PAGE and immunoblotting

Cells were placed on ice and rinsed twice with cold phosphate buffered saline (PBS) (Wisent) and collected for lysis by scraping. Collected cell suspensions were centrifuged at 4°C and 200 x g. Supernatant was removed, cell pellets were lysed in RIPA lysis buffer [50 mM Tris-HCl pH 7.5, 150 mM NaCl, 0.5% sodium deoxycholate, 1 mM PMSF, 1 mM DTT, 1% NP40, 0.1% SDS, supplemented with 1X protease inhibitor (Roche) and 1x PhosSTOP (Roche)] and incubated on ice for 15 min with occasional vortexing. Lysates were cleared by centrifugation at 4°C 13,000 x g for 15 min and protein-containing supernatants moved to new 1.5 ml centrifuge tubes. Protein quantification was done by Pierce BCA kit (ThermoFisher Scientific), followed by equalization of concentration and preparation for SDS-PAGE. Samples were then transferred to nitrocellulose membranes (Cytiva) for immunoblotting. Membranes were blocked for 1 h at room temperature in 3% bovine serum albumin (BSA) (BioBasic) or 3% nonfat dry milk powder, as instructed by antibody manufacturers. Incubation with primary antibodies was done overnight at 4°C with constant agitation, while secondary antibody incubation was done for 1 h at room temperature. Tris-buffered saline (TBS), supplemented with Tween (BioBasic) at 0.1 % to obtain TBST, was used to rinse membranes in three 10 min intervals, after each antibody incubation. One final 10 min TBS wash followed before chemiluminescent detection using substrate solution (BioRad). The following antibodies were used in the study:

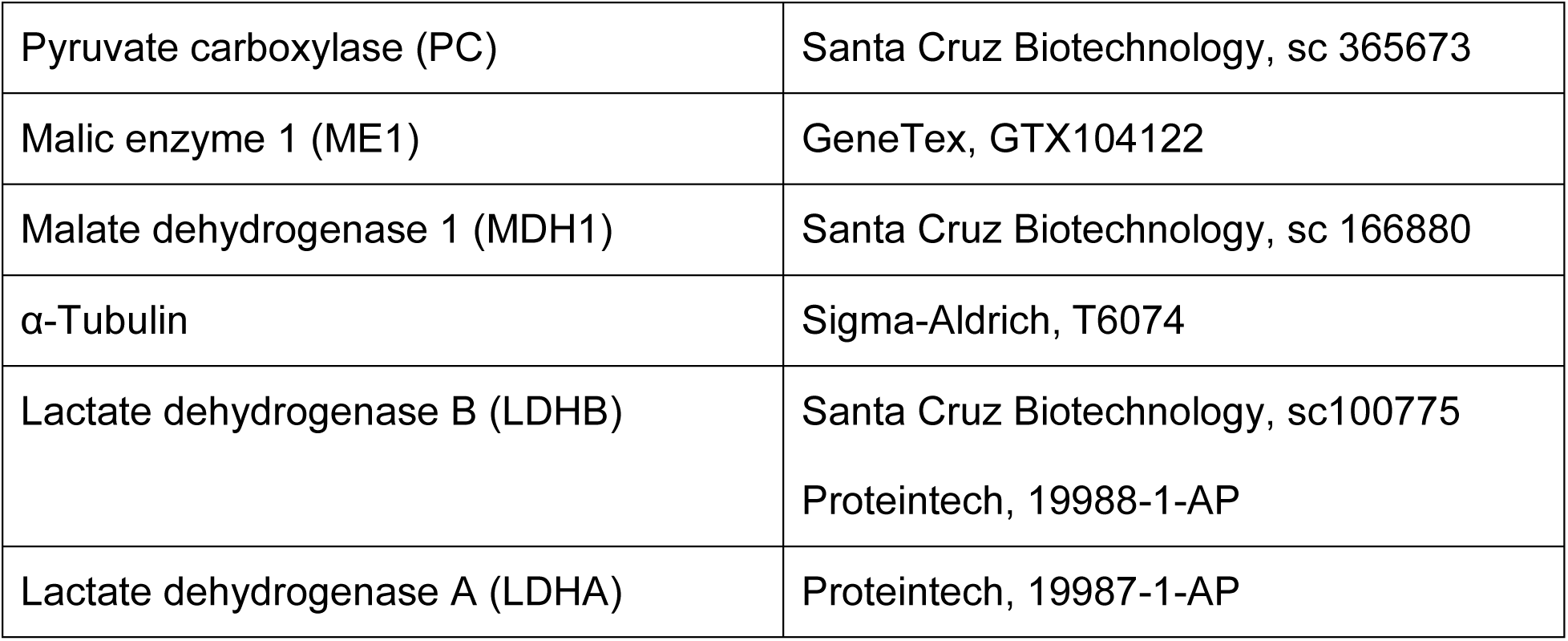

|  |  |
| --- | --- |
| Pyruvate carboxylase (PC) | Santa Cruz Biotechnology, sc 365673 |
| Malic enzyme 1 (ME1) | GeneTex, GTX104122 |
| Malate dehydrogenase 1 (MDH1) | Santa Cruz Biotechnology, sc 166880 |
| α-Tubulin | Sigma-Aldrich, T6074 |
| Lactate dehydrogenase B (LDHB) | Santa Cruz Biotechnology, sc100775<br>Proteintech, 19988-1-AP |
| Lactate dehydrogenase A (LDHA) | Proteintech, 19987-1-AP |

### Cell proliferation assay

Cell proliferation was measured by determining live cell numbers at indicated timepoints. Live cell counts were obtained through trypan blue exclusion using the Countess automated cell counter (Thermo Fisher Scientific). Alternatively, when indicated, crystal violet (Bioshop) retention assay was used as described previously(Igelmann *et al*., 2021).

### Xenograft experiments

5 x 10^4^ breast cancer cells were injected within the fourth mammary fat pad (orthotopically). Cells were resuspended after two PBS washes in ice-cold PBS and mixed in a ratio of 1:1 with matrigel (Corning) on ice. Mixture volumes of 50 µL were injected using 0.5 mL syringes. Female 6-8-week-old mice were matched by background to breast cancer cell line used for the experiment (SCID-Beige for NMuMG-NT and BALB/c for 67NR, 4T1-526 and 4T1). Formed tumors were monitored and measured with calipers using the following formula: volume = 4/3 × (3.14159) × (length/2) × (width/2)^2^.

### Spontaneous metastasis assay

All metastases were derived from orthotopically injected tumors. Lung metastases were examined upon animal necropsy, as lung tissue was formalin-fixed, paraffin embedded, sectioned and hematoxylin and eosin (H&E) stained. Necropsies were performed either as tumor volumes reached 1000 mm^3^ or 21 days after resecting tumors at a volume of 300 mm^3^. Stained slides were then scanned (Aperio 20X), classified as metastatic or non-metastatic, and metastatic area quantified using ImageScope.

The limitations of the metastatic models were considered when designing spontaneous metastasis experiments, such that genetic alterations expected to increase metastasis were introduced in two models with lower metastatic ability (e.g., 67NR, NMuMG-NT), while genetic approaches hypothesized to decrease metastasis were employed in models with higher metastatic capacity (e.g., 4T1, 4T1-526). To avoid confounding of the differences in primary tumor growth between the conditions, resections or necropsies took place at equal primary tumor volumes. Controlling for tumor volume necessarily leads to altered lengths of time from tumor implantation to lung collection. However, in each case faster-growing tumors also formed more metastasis, despite a shorter time from orthotopic injection to lung collection. Consequently, this may leave room for an underestimation of actual fold changes of metastasis within the examined groups. Lastly, metastases were quantified as a percentage of the total lung area covered by metastatic lesions. When such analyses were precluded by either complete lack of metastases in certain groups, or unclear borders between metastatic and non-metastatic areas, lungs were only designated as “metastatic” or “non-metastatic” and shown as percentages.

### Co-immunoprecipitation

MDA-MB-231 and/or MDA-MB-468 cells were collected at 80% confluency from two 15 cm cell culture dishes after being washed twice with ice-cold PBS. Cells were scraped into 1 mL of immunoprecipitation (IP) buffer (50 mM Tris-HCl pH 7.9, 1 mM EDTA, 0.1 mM EGTA, 12.5 mM MgCl_2_, 400 mM NaCl, 20% glycerol, 1% Triton X-100, 0.1% SDS and 1X EDTA-free protease inhibitor cocktail (Roche). Cell lysates were sonicated for 30 s and then cleared at 16,300 x g for 30 s. For input controls, 1% of the lysate was taken. Immunoprecipitation was done using 5 μg of anti-ME1 rabbit polyclonal antibody (Genentex), or rabbit pre-immune serum (Cell Signaling Technologies) overnight at 4°C with end-over-end rotation. Recovery of immunoprecipitated proteins was performed using protein G Dynabeads (Thermo Fisher), which were incubated with the lysates for 2 h and then washed twice for 10 min and three times for 60 min at 4°C with IP buffer. Proteins were eluted by incubation at 95°C in 6X SDS loading buffer for 5 min, with occasional vortexing.

### ME1 immunoprecipitation & mass spectrometry analysis

ME1 immunoprecipitation was done as described in section “Co-immunoprecipitation” but was done in PC-3 cells cultured to 80% confluency in 15 cm plates. Modifications to protocol were done and include two additional 1 h washes followed by overnight incubation with wash buffer containing 500 mM NaCl. Samples were then incubated with magnetic beads against protein A-G (Thermo Fisher). Beads containing immunoprecipitate were then washed twice for 30 min in wash buffer. After the last wash, beads were processed for mass spectrometry (MS). For each sample, released proteins were loaded onto a single stacking gel band to remove lipids, detergents and salts. The gel band was reduced with DTT, alkylated with iodoacetic acid and digested with trypsin. Extracted peptides were re-solubilized in 0.1% aqueous formic acid and loaded onto a Thermo Acclaim Pepmap (Thermo, 75 µM i.d. x 2 cm C18 3 µM beads) precolumn and then onto an Acclaim Pepmap Easyspray (Thermo, 75 µM x 15 cm with 2 µM C18 beads) analytical column separation using a Dionex Ultimate 3000 uHPLC at 250 nL/min with a gradient of 2-35% organic (0.1% formic acid in acetonitrile) over 3 hours. Peptides were analyzed using a Thermo Orbitrap Fusion mass spectrometer operating at 120,000 resolution (FWHM in MS1) with HCD sequencing (15,000 resolution) at top speed for all peptides with a charge of 2+ or greater. The raw data were converted into *.mgf format (Mascot generic format) for searching using the Mascot 2.6.2 search engine (Matrix Science) against Mouse proteins (Uniprot 2022). The database search results were loaded onto Scaffold Q+ Scaffold_5.0.1 (Proteome Sciences) for statistical treatment and data visualization. After removal of creatine and other contaminants, clusters of identified putative-interacting protein targets were displayed using STRING analysis.

### Ex vivo tumor metabolic labelling

SCID-Beige mice were orthotopically injected with 5 x 10^4^ NMuMG-NT cells and tumors grown to a volume of 300 mm^3^. Tumor resection was then performed in sterile conditions, under a biological safety cabinet, while animals were anesthetized with isoflurane. Excised tumors were immediately put into cold transfer media [DMEM: HAM’s F12 = 2:1, FCS 2% (Wisent), hydrocortisone 0.3 μg/ml (Sigma), 1x ITS (Wisent), 3,3’,5 Triiodothyronine 1 ng/ml (Sigma), EGF 8 ng/ml (Sigma), cholera toxin 7 ng/ml (Sigma), adenine 0.2 mg/ml (Sigma), 1% penicillin/streptomycin (Wisent), 20 mM HEPES (Wisent)] and further processed for wet cutting at the vibratome (removal of excess fat, removal of large tendons and peritoneal fascia, in case of peritoneal invasion). Tumors were stored for a maximum of 4 h in 4°C transfer medium.

To section tumor tissue using a vibratome, tumors were embedded in 4% low-melt agarose (Biobasic) dissolved in PBS (Wisent). To embed the tissue, it was deposited on a cushion of 1 mL of low melt agarose which was cooled to 37 °C. Plastic molds were immediately transferred on ice until hardening. They were then removed and the remaining agarose block was glued with Krazy glue. The vibratome chamber was filled with PBS and exterior chamber was filled with ice. Cuts of 1 mm were initially done to reveal the tissue, after which step cuts of 300 µm were performed. Settings of the vibratome were the following: 0.60 mm/s speed, 1.20 mm amplitude, 300 µm auto feed. Tissues were recovered from PBS with a paint brush and transferred into a 6-well plate with appropriate NMuMG-NT cell culture growth media. Tumor tissue slices were incubated for a minimum of 16 h at 37 °C in a humidified CO_2_ incubator with constant shaking at 30 RPM.

To prepare samples for mass spectrometry slices were transferred with the brush into a new 6-well plate and incubated for the appropriate time in the presence of required isotope-labelled metabolites. Tissues were washed twice in ice-cold 150 mM pH 7.4 ammonium formate and transferred into a 2 mL tube filled with 50% MeOH/H20 or 80% MeOH/H20 and 4 beads. Tubes were beaten in bead beater for 2 min at 30 Hz. Process was repeated for a minimum of 3 times.

### [2-^2^H] Lactate tracing

5 x 10^5^ cells were seeded 16 h prior to the experiment. On the day of experiment the media was removed and replaced with fresh DMEM containing 5 mM of unlabelled lactate and incubated for 2 h. The equilibration media was replaced with DMEM containing 5 mM of labelled sodium lactate (Sodium L-lactate (3,3,3-D₃, 98%) 20% w/w in water, Cambridge Isotope Laboratories) and incubated for 6h. The tracing was performed as described before(Igelmann *et al*., 2021). Briefly, media was removed, and cells washed three times with isotonic saline solution at 4°C on ice and quenched with 600 μL of 80% methanol on dry ice. Cells were detached by scraping from the plates and transferred to prechilled tubes. Cell suspensions were sonicated for 10 min at 4°C in cycles of 30 sec ON, 30 sec OFF, at high setting with Diagenode Bioruptor. Sonication was repeated to ensure complete recovery of metabolites. Cell debris were removed through centrifugation (16,000 x g, 4°C). Supernatants were transferred to prechilled tubes containing 1 µL of 750 ng/μL ^2^H_27_-myristic acid and dried by vacuum centrifugation (Labconco) with sample temperature maintained at 4°C overnight. Dried pellets were dissolved in 30 μL of pyridine containing methoxyamine-HCl (10 mg/mL) using a sonicator and vortex. Samples were then spun down at 16,000 x g, incubated for 30 min at 70°C and then transferred to GC-MS injection vials containing 70 μL of N-tert-Butyldimethylsilyl-N-methyltrifluoroacetamide (Sigma). Sample mixtures were further incubated at 70°C for 1 h. Out of each sample, 1 μL was injected for GC–MS analysis. GC–MS instrumentation and software were all from Agilent. GC–MS methods and mass isotopomer distribution analyses were done as previously described (Gravel *et al*, 2016). Briefly, GC-MS analysis was performed on an Agilent 5975C series GC-MSD with triple-axis HED/EM detector, equipped with DB-5MS + DG capillary column (30 m length, 10 m Duraguard deactivated fused silica tubing, 0.25 mm internal diameter, 0.25 μm film thickness) (Agilent) and a 7890A gas chromatograph, with a 7693 autosampler. Out of each sample 1 μL was injected into the GC in splitless mode with inlet temperature at 280°C with helium gas as the carrier at a flow rate at 1.552 mL/min. The quadrupole was set to 150°C and the GC-MS interface at 285°C. The oven program for all metabolite analyses started at 60°C, held at 1 min and increased at a rate of 10°C/min up to 320°C. Bake-out was at 320°C for 9 min. Ions were generated by Electron Impact (EI) set at 70eV. Sample data were acquired both in scan (1-600 m/z) and selected ion monitoring (SIM) modes.

Metabolites were identified by matching mass spectra and retention times using authentic standards (Sigma). Metabolites were quantified by the integration of the M-57 ion response divided by the response of the internal standard ^2^H_27_-myristic acid and normalized to cell count. For stable isotope tracer analysis, the integration values for quantifier (m+0) and all possible isotopomers (m+1, m+2, etc.) are summed up to obtain the total amount of metabolite. The mass isotopomer distribution (MID) vector is obtained by dividing the m+0 and all isotopomer by the total amount of metabolite. The corrected MID is obtained by removing the contribution of naturally occurring stable isotopes (McGuirk *et al*, 2013).

### Gene overexpression and rescue via lentiviral transduction

Lentiviral transductions were done by transfecting HEK293T cells with 12 µg of target vector, 8 µg psPAX2 and 4 µg pMD2.G. Transfections were performed using Jetprime (Polyplus), in 10 cm dishes when cultures reached 60% confluency, according to the manufacturer’s instructions. Growth medium was replenished 24 h after transfection, and virus collected the following three days. Each day of virus collection, recipient cells were transduced in 6-well plates with a mix of filtered (0.45 µm, FroggaBio) viral soup and full growth media (1:1) and supplemented with 8 µg/mL polybrene (Sigma Aldrich). Cells expressing selection markers (i.e., fluorescent proteins) were collected through fluorescence-activated cell sorting (FACS) into 6 well dishes, to be expanded and used for downstream analysis.

### Gene depletion mediated by CRISPR-Cas9

In order to deplete the endogenous expression of malic enzyme 1 (ME1), malate dehydrogenase 1 (MDH1), and pyruvate carboxylase (PC) in 4T1-526 cells, two guides (sgRNAs) were designed against each gene using CHOPCHOP (https://chopchop.cbu.uib.no/) and purchased with the appropriate overhangs to allow cloning into the lentiCRISPRv2 vector expressing neomycin (G418) resistance, hygromycin B resistance, or green fluorescent protein (GFP), respectively, alongside Cas9 and a gRNA cloning site (Addgene #98292, #98291, #82416). Cloning was carried out as previously described (Sanjana *et al*, 2014; Shalem *et al*, 2014), with the following modifications: *i)* using BsmBI-v2 (New England Biolabs) to digest the lentiCRISPRv2 plasmids, *ii)* using the Zymoclean Gel DNA Recovery Kit (Zymo Research), and *iv)* carrying out the ligation reaction overnight at 4°C. Transformation of the ligated plasmid was carried out through a 42°C heat-shock into chemically competent Stbl3 cells (Fisher Scientific). Bacteria were plated on Lysogeny Broth (LB)-agar plates (Biobasic) containing ampicillin (1 µg/mL) (BioBasic) and grown colonies were picked and inoculated into Lysogeny Broth with ampicillin (100 μg/mL) overnight at 37°C. Extraction of plasmid DNA was done using the QIAprep Spin Miniprep Kit (Qiagen) and samples were sent for Sanger sequencing using a primer for the human U6 promoter (GAGGGCCTATTTCCCATGATT). Correct insertion of the gRNA into the plasmid was verified before proceeding.

For lentiviral production, HEK293T cells were seeded in 6 cm dishes to achieve 60% confluency the following day. Then, cells were transfected with 4 µg of lentiCRISPRv2 plasmid containing one of the gRNA inserts targeting each of the three candidate genes (resulting in 6 different plates, since each gene was targeted by two guides), or empty-lentiCRISPRv2 plasmid. Simultaneously, 2.66 µg of psPAX2 packaging plasmid (Addgene #35002), and 1.66 µg of pMD2.G plasmid (Addgene #12259) was added, and transfection was done using the jetPRIME transfection reagent (Polyplus). Growth media was changed 24 h later and collected 48 h post-transfection. The virus-containing media from the various 6 cm plates was filtered through a 0.45 µm filter (Frogga Bio) and was mixed with fresh media at a 1:1 ratio. The virus media targeting ME1, MDH1 and PC were combined, as were the virus media that contained empty vectors. Polybrene (Sigma-Aldrich) was added to obtain a final concentration of 8 µg/mL. The mixture was added to 1.0×10^4^ 4T1-526 cells that were seeded in 6-well plates. Cells were re-transduced the following two days, following the same procedure. All cell lines were allowed to recover for 48 h after the last transduction. Then, cell lines were selected with 500 µg/mL G418 and 1 mg/mL hygromycin B for up to 7 days and allowed to recover for an additional three days prior to fluorescence-activated cell sorting (FACS). Single cell sorting for high GFP expressing cells was conducted using a BD FACSAria Fusion Flow Cytometer (BD Biosciences). Cells expressing GFP were selected and grown in 96 well plates containing conditioned media (45% filtered culture media, 45% fresh media, 10% FBS). After approximately two weeks, sorted clones were transferred to larger plates.

Depletion of endogenous ME1, MDH1, and PC protein levels in 4T1-526 cells was initially confirmed by immunoblotting. Candidate clones with loss of ME1, MDH1, PC expression were kept for further validation. Final validation of gene deletion was done through genomic DNA extraction using PureLink Genomic DNA Mini Kit (Thermo Fisher Scientific) and subsequent Sanger sequencing using primers flanking the regions of interest.

### Transwell migration/invasion assay

Cells were washed twice with PBS and cultured in medium devoid of serum for 16h. Then, cells were detached by trypsinization (Wisent), neutralized in full growth media and washed twice with PBS by alternating centrifugation and resuspension. Next, cells were resuspended in serum-devoid medium and seeded in to transwell migration inserts (12-well format, 8 µm pore size, Corning) in 1 mL volumes for cell migration assays. For cell invasion assays, the inserts were previously coated with 300 µL of a 300 µg/mL suspension of matrigel (Corning) in PBS for ∼60 min at +37°C, 5% CO_2_. Full growth medium was placed in the bottom chamber at a volume of 2 mL to initiate migration/invasion of cells. After 24h, inserts were collected, media aspirated, and non-migrated or non-invaded cells removed from the inner side of the insert with a cotton-tip swab, leaving only the migrated or invaded cells on the outer side of the insert. These cells were then fixed for 20 min in 10% formalin and stained with 0.1% crystal violet for 20 min. After rinsing with water and drying for 24h, the inserts were imaged and numbers of migrated/invaded cells determined.

### Anoikis resistance assay via Annexin Vand DAPI analysis

Anoikis resistance assays were performed using the APC Annexin V Kit (Biolegend) according to manufacturer’s instructions. Cells were seeded at 5 x 10^5^ cells on either regular cell culture plates, or non-treated plates (Fisher Scientific) coated for 5 min with Anti-Adherence Rinsing Solution (Stem Cell Technologies) and then rinsed. Cells were then incubated for 24 h. Subsequently, suspended cells were collected and spun for 10 min at 350 x g. Pellets containing dead and viable cells were washed twice with cold Cell Staining Buffer (Biolegend) and then resuspended in Cell Staining Buffer. APC Annexin V Biolegend 640920 (dilution 1:50) and DAPI (Invitrogen) at 300 ng/mL were added and incubated for 15 min at room temperature in the dark. The reaction was quenched by adding 400 μL of BioLegend Cell Staining Buffer. Samples were then passed through a 35 μm cell strainer into glass round-bottom tubes (Falcon) and analyzed using flow cytometry (BD Fortessa). Final analysis was done using Flow JO.

### RNA extraction

Prior to isolation of RNA, all contact surfaces were cleaned using RNaseZAP (ThermoFisher Scientific) and precaution was taken to maintain RNase-free conditions. Extraction of RNA was performed using TRIzol reagent (Thermo Fisher Scientific), according to manufacturer’s instructions. After fully resuspending cell cultures in 1000 µL of TRIzol, 200 µL of chloroform (Sigma Aldrich) was added and tubes mixed vigorously to homogenize. Aqueous phase was isolated after 15 min of centrifugation at 13,000 x g in +4°C. RNA was precipitated with 2X volume of isopropanol (Bio Basic), supplemented with 2 µL of Glycoblue reagent (Thermo Fisher Scientific), and incubated at -80°C for 30 min. Next, a 20 min centrifugation at 13,000 x g in +4°C followed. Supernatant was carefully removed to reveal RNA pellets, which were then washed twice in 1000 µL of cold 75% ethanol. Supernatant was removed once more and RNA pellets left to dry (∼10 min or until translucent). RNA was resuspended in 40-100 µL of sterile molecular grade H_2_O (Wisent). RNA was stored at -80°C until further use.

### RTqPCR analysis

Total extracted RNA was used to synthesize complementary DNA (cDNA) using SensiFAST cDNA synthesis kit (FroggaBio), whereby 500 ng of template was used. Quantitative polymerase chain reaction (qPCR) was performed on BioRad systems using SensiFAST SYBR Lo-Rox Mix (FroggaBio). All steps were done according to manufacturer’s instruction. The following primers were used for RTqPCR analysis:

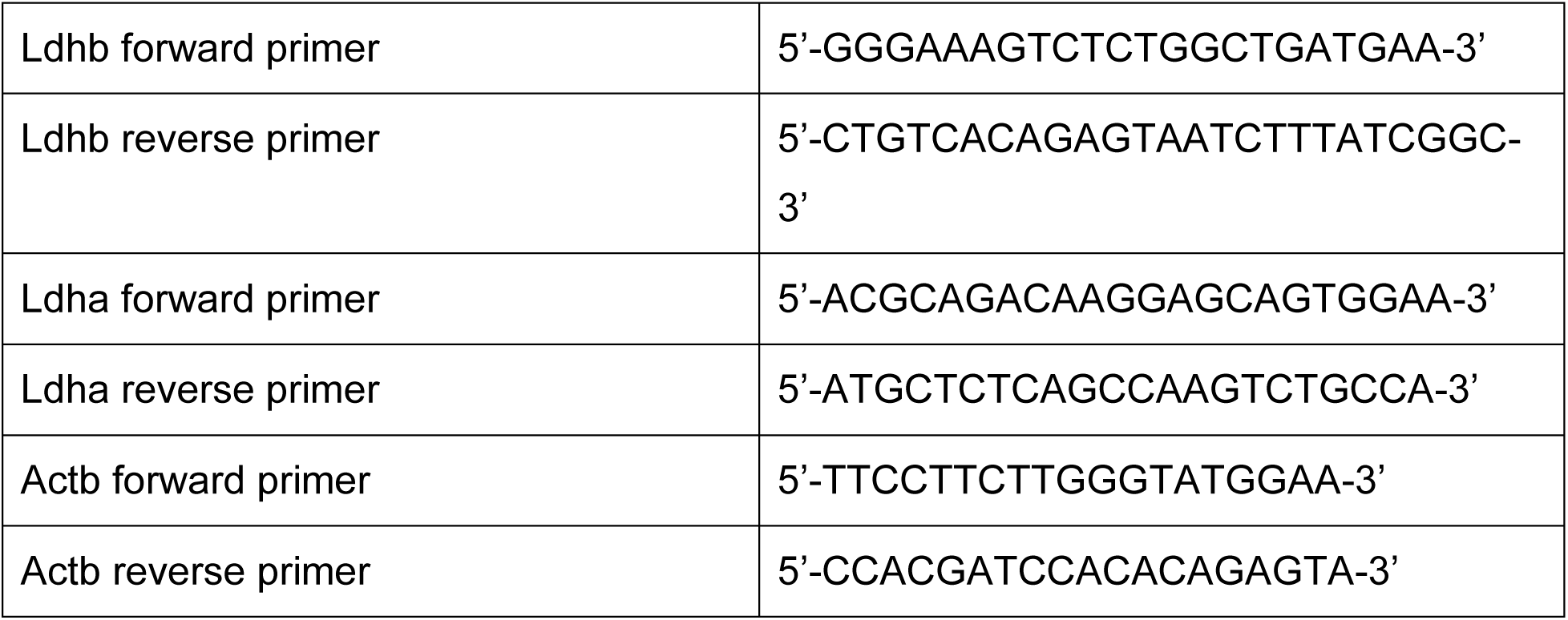

## Acknowledgements

We would like to thank Christian Young for assistance with flow cytometry experiments and Frédérick A. Mallette for generously gifting us plasmids encoding IDH1. Funding sources: Terry Fox Foundation Oncometabolism Team Grant TFF-242122, Canadian Institutes for Health Research PJT-183843 (to I.T), PJT-451236 (to I.T. and G.F), and PJT-203988 (J.U-S). Canadian Cancer Society (CCS) and Lotte & John Hecht Memorial Foundation Disruptive Innovation Grant in Cancer Research (DI-25) #708583 (to G.F.). S.I. received funding from EMBO postdoctoral fellow (ALTF 13-2022) and from Research Foundation Flanders (FWO) junior postdoctoral fellowship (1266225N). P.J. is supported by a Fonds de recherche du Québec (FRQ-S) Doctoral Training award. D.P. is supported by Cancer Research Society Next Generation of Scientists Award. N.C. is supported by the Peter Quinlan Fellowship in Oncology (Faculty of Medicine and Health Sciences, McGill University). I.T. is the Canada Research Chair in Regulation of mRNA Translation and Metabolism. G.F. is the CIBC Chair for Breast Cancer Research. Metabolic analysis was performed at The Metabolomics Innovation Resource (MIR), McGill University supported by the Canada Foundation for Innovation, The Dr. John R. and Clara M. Fraser Memorial Trust, the Terry Fox Foundation (TFF-242122) and McGill University.

## Author contributions

Conceptualization: IT, GF, JUS, SI, PJ

Experimental work: SI, PJ, VB, NC, LZ, RP, DP, SM, TNG, LT

Data analysis/interpretation: SI, PJ, VB, NC, LZ, DP, DA, MP, TNG, LT, JFT, JUS, GF, IT

Funding acquisition: IT, GF, JUS

Supervision: IT, GF, JUS

Writing – original draft: PJ, IT

Writing – review & editing: IT, GF, JUS, SI, PJ

